# Near-Optimal P300 Speller Performance Using Large Language Models: A Multi-Model Analysis with Performance Bounds

**DOI:** 10.1101/2025.10.28.685216

**Authors:** Nithin Parthasarathy, James Soetedjo, Saarang Panchavati, Nitya Parthasarathy, Dongwoo Lee, Corey Arnold, Nader Pouratian, William Speier

## Abstract

Amyotrophic lateral sclerosis (ALS), a progressive neurodegenerative disease, severely impairs communication, requiring assistive technologies that restore interaction. The P300 speller brain-computer interface (BCI) enables communication by translating EEG responses into text; however, its practical adoption is limited by slow typing speed and the need for subject-specific calibration. Recent work has demonstrated that large language models (LLMs), such as GPT-2, can significantly improve the performance of the P300 speller by predicting words. However, it remains unclear whether these gains are model-specific or represent a broader trend across language models. Furthermore, the fundamental performance limits of LLM-assisted P300 spellers have not been systematically characterized within a unified decoding framework. In this study, we address these gaps through a systematic multi-model theoretical analysis framework. We evaluate a wide range of language models and introduce an idealized LLM to establish upper bounds on achievable performance. In addition, we incorporate cross-subject classifier training to reduce calibration requirements and assess generalization across subjects. Using extensive simulations on EEG data from 78 subjects, we demonstrate that the evaluated models consistently achieve substantial improvements in typing speed, with gains of up to ∼40% (across-subject training) and ∼75% (within-subject training) over conventional approaches. More importantly, we show that multiple models, despite architectural differences, operate within 5% of the theoretical performance bound, indicating diminishing returns from further model scaling. These improvements generalize across both within-subject and across-subject classifiers. Our results suggest that LLM-assisted P300 spellers are approaching their fundamental performance limits within the considered decoding framework, shifting the primary bottleneck from language modeling to neural signal decoding. This work provides both a practical framework for improving BCI communication and a theoretical perspective on its achievable limits.

## 1. Introduction

Amyotrophic lateral sclerosis (ALS) is a progressive neurodegenerative disease that severely impairs communication [1]. Brain-computer interfaces (BCIs) [2]-[7], and in particular the P300 speller, provide a non-invasive means of restoring communication by translating neural responses into text. In the P300 speller, a character matrix—referred to here as a *flashboard* —is presented to the subject, and the resulting EEG response to highlighted characters is recorded, processed, and interpreted [3, 7]. Despite its clinical promise, practical deployment of P300 spellers remains limited by low communication speed and the need for subject-specific calibration [5].

A promising direction to address these limitations is the integration of natural language processing (NLP) into the P300 decoding pipeline. Early work from our group and others demonstrated that character-level and n-gram language models can improve speller performance by exploiting the statistical structure of natural language [6, 9, 10, 14, 15]. However, these approaches treat character selections as largely independent events and provide limited gains from context beyond a few preceding characters. More recently, we demonstrated that large language models (LLMs)—specifically GPT-2—can substantially improve P300 speller performance through word-level prediction, and that these gains generalize across subjects via cross-subject classifier training [12, 13]. That study established GPT-2 as a strong baseline and introduced the cross-subject evaluation framework (ASCV) used here. However, two fundamental questions were left unanswered: (i) whether these gains are specific to GPT-2 or represent a general property of language models broadly, and (ii) what the fundamental performance limits of LLM-assisted P300 spellers are. The current study directly addresses these gaps.

To this end, we evaluate a broad range of open-source language models spanning encoder- and decoder-based architectures of varying scales, from encoder-based models such as RoBERTa [21] and BERT [22] to decoder-based models such as GPT-2 [34], LLaMA [32, 33], Falcon [30], and others [23, 24, 25, 26, 27, 28, 29, 31]. We introduce a formal theoretical framework—oracle performance bounds and a Shannon physiological ceiling—to establish how close practical systems are to the fundamental performance limits of this LLM-assisted P300 speller architecture. Together, these contributions provide a system-level characterization of performance limits that cannot be inferred from single-model studies.

The central objective of this study is to characterize the performance limits of LLM-assisted P300 spellers. To this end, we introduce an idealized LLM capable of perfect word prediction to establish an upper bound on achievable performance. This bound is then used to evaluate how closely these models approach optimal operation. In addition, we investigate the role of cross-subject classifier training—introduced in our prior work [13]—in reducing or potentially eliminating the need for subject-specific calibration, and assess whether performance gains generalize across subjects and model architectures.

As will be shown, many of the evaluated models operate within a small margin (*∼*5%) of this bound, indicating that performance may be approaching fundamental limits. Furthermore, this performance substantially exceeds that achievable using character-level prediction alone. While the experimental framework, dataset, and decoder structure follow our prior work [13], the present study introduces three key extensions. First, we evaluate a broad range of modern large language models across architectures and scales, rather than a single model. Second, we introduce formal oracle-based performance bounds to quantify the achievable limits of language-model-assisted decoding. Third, we identify and characterize a convergence phenomenon across language model architectures, leading to a system-level interpretation of performance limits. Together, these contributions provide a new system-level perspective on the performance limits of P300 speller systems. Our results suggest that further improvements in language modeling alone are unlikely to yield significant gains, shifting the focus of P300 speller research toward neural decoding and signal acquisition.

Our technique involves extensive *offline* simulations using multi-subject EEG data, incorporating layered word completion algorithms (e.g., GPT-2, LLaMA) in conjunction with Dijkstra’s algorithm. This framework enables systematic comparison of performance across models and classifier configurations, including both *within*-subject and *across*-subject training. Additional system-level optimizations, including flash-board design and scanning strategies, are incorporated to assess their interaction with language-model-based decoding.

The simulation techniques developed in this work allow for the comparison of many speller scenarios on large and representative targets. These include OOV words that would otherwise be infeasible in an online study due to the significant time requirements. Evaluation of these findings in online settings will be the focus of future work.

## 2. Contributions

Specifically, this work makes the following key contributions:

- **Multi-model generalization analysis:** We evaluate a broad set of modern large language models across different architectures and scales, demonstrating that performance gains in P300 spellers are largely model-agnostic.
- **Theoretical performance bounds:** We introduce an idealized language model to establish upper bounds on achievable performance, enabling quantitative assessment of how closely practical systems approach optimal operation.
- **Convergence of model performance:** We show that all evaluated models operate within 2–5% of the theoretical bound, revealing a previously unreported convergence phenomenon and indicating diminishing returns from further model scaling.
- **Bottleneck identification:** We demonstrate that the primary limitation in modern P300 spellers has shifted from language modeling to neural signal decoding and physiological constraints.
- **Cross-subject validation:** We show that these results generalize across both within-subject and across-subject training paradigms, supporting the feasibility of reducing or potentially eliminating subject-specific calibration.

The organization of this paper is as follows. In Section 3, a detailed description is provided of the various techniques and algorithms used for the flashboard, stimuli, and classifiers, as well as character and word prediction. Section 4 illustrates the results, while Section 5 provides a summary of the results, which is followed by the section on conclusions and future directions.

## 3. Methods

The methods employed in this study follow the framework established in our prior work [13]. We summarize each component below, noting where the current study introduces new elements or departs from the prior approach.

### 3.1. Language models

This section describes the multi-level character language model used for flashboard probability estimation and out-of-vocabulary (OOV) handling, following the approach of [13]. Character frequency in English motivates the scanning order: vowels occur more frequently than consonants [20], and are therefore prioritized earlier in the flashboard sequence. Conversely, less frequent consonants, such as *‘x’* and *‘z’*, are highlighted later.

Smoothing techniques [16] enable transitions across models when words are encountered outside the vocabulary (OOV), blending higher-order and lower-order models. As illustrated in Figure 2, the language model spans a hierarchy ranging from simple trigram models to more complex biword models.

**Figure 1:**
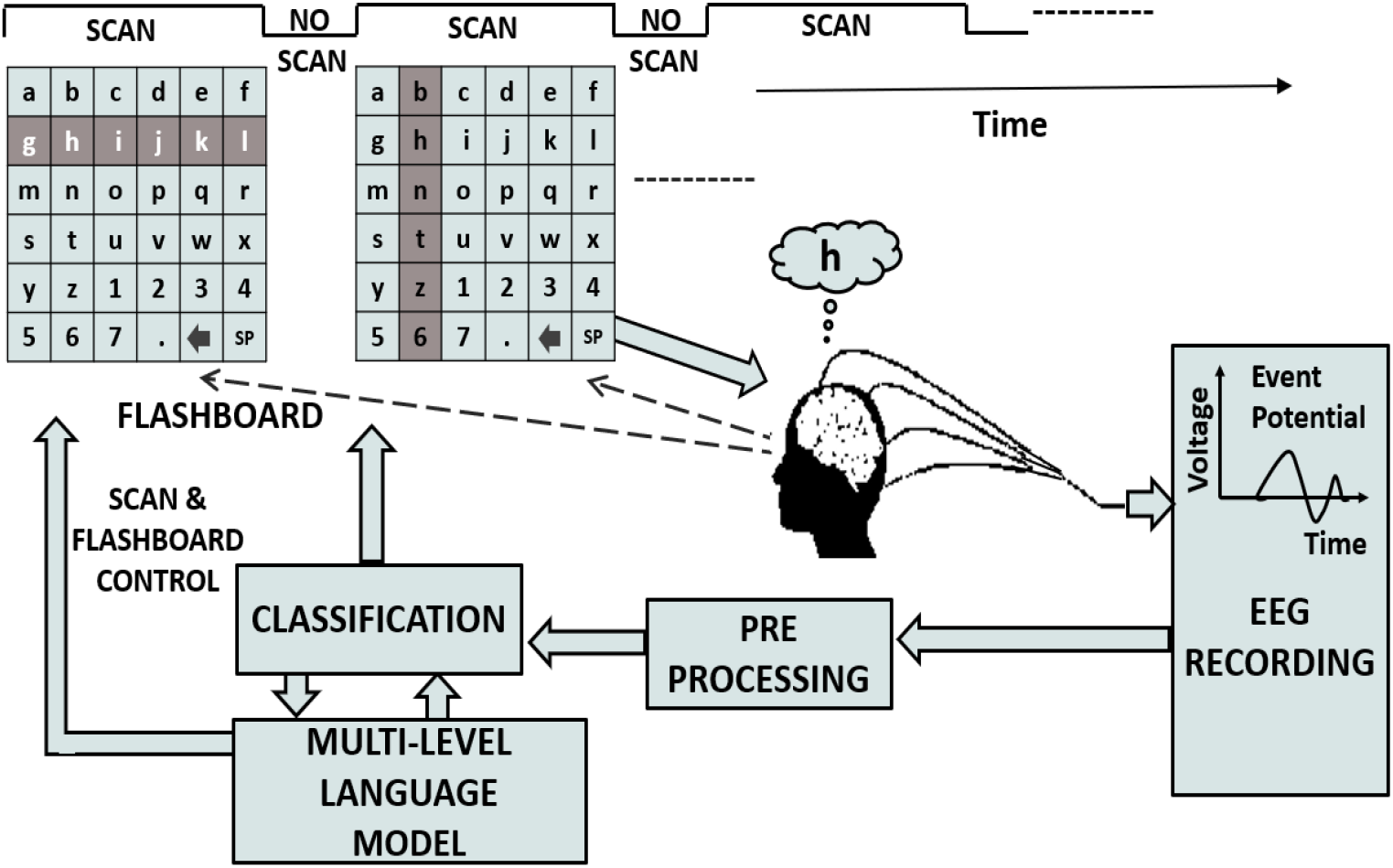
A block diagram of the P300 speller based BCI used in this work. Subject EEG response to character highlights on a flashboard are shown.

**Figure 2:**
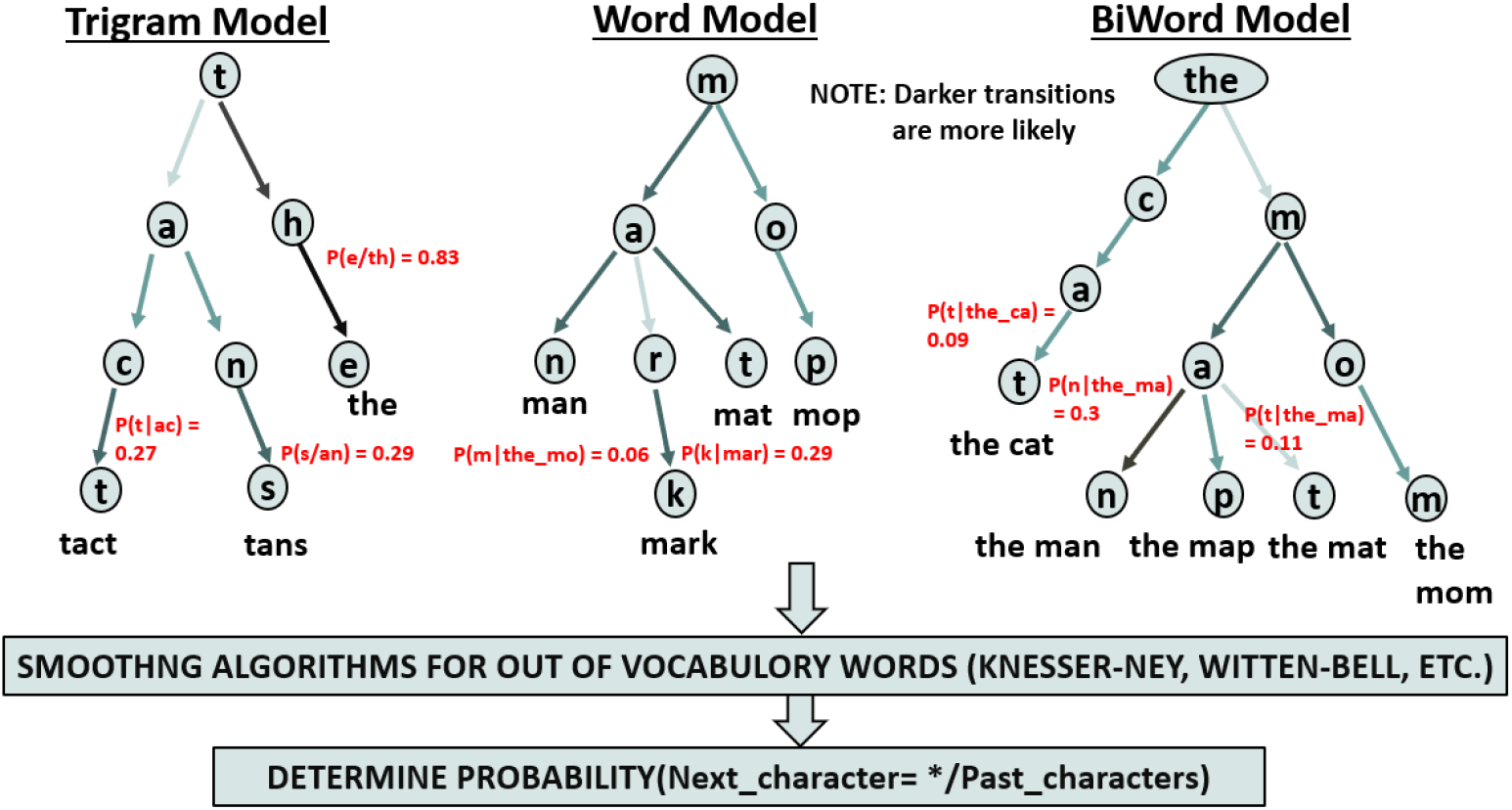
Types of language models used. Starting with the simplest on the left to the most complex on the right

As an example, consider the word *‘cat’* and suppose *‘ca’* has already been spelled. If *‘cat’* is not present in the language model, the conditional probability *p*(‘*t*’ |‘*ca*’) is zero. Smoothing addresses this by using alternative contexts, such as *p*(‘*t*’ |‘*a*’), where characters have been observed to co-occur.

Among several smoothing techniques evaluated, Kneser–Ney smoothing was selected for its strong OOV performance [37]. This approach recursively combines higher-order and lower-order n-gram models to produce robust probability estimates.

The frequency counts used in the language model were estimated from the Brown Corpus [17], a widely used benchmark dataset that provides broad coverage of written American English. The complete recursive formulation of the smoothing equations is provided in Appendix A.

### 3.2. Data Collection

The dataset used here is identical to that described in our prior work [13]; we summarize the key parameters for completeness. Data for offline analyses were obtained from 78 healthy volunteers with normal or corrected-to-normal vision between the ages of 20 and 35. Figure 3 shows the BCI display flashing before a volunteer whose response is being recorded. All data were acquired using GTEC amplifiers, active EEG electrodes and electrode cap (Guger Technologies, Graz, Austria); sampled at *256 Hz*, referenced to the left ear; grounded to *AF·Z*; and filtered using a bandpass of *0*.*1* – *60 Hz*. The set of electrodes consisted of 32 channels placed according to a previously published configuration [36]. We used a window of 600 ms at 256 Hz, resulting in 154 time points per channel. We then downsampled the signal by a factor of 12 to obtain a feature vector that was 32×13=416 features. The system also used a *6* × *6* character grid, row and column flashes, and a stimulus duration of 100 msec and an interval between stimulus of 25 ms for a stimulus onset asynchrony of 125 msec. The channels used were *Fpz, Fz, FC1, FCz, FC2, FC4, FC6, C4, C6, CP4, CP6, FC3, FC5, C3, C5, CP3, CP5, CP1, P1, Cz, CPz, Pz, POz, CP2, P2, PO7, PO3, O1, Oz, O2, PO4, PO8* again based on a published montage [36].

**Figure 3:**
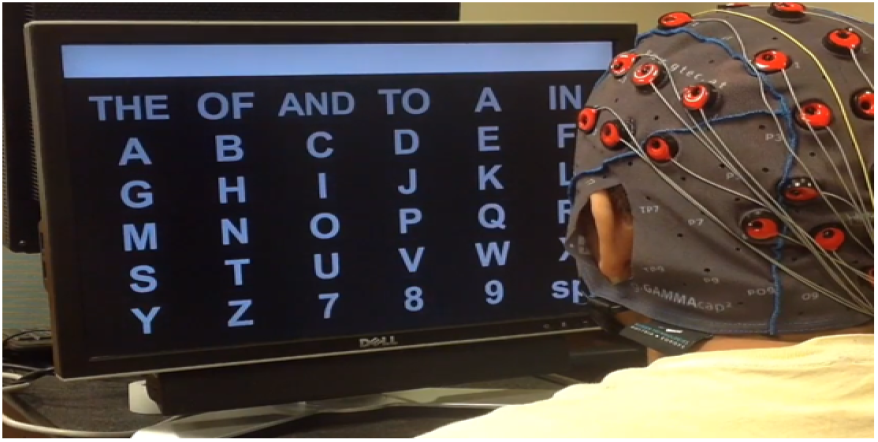
BCI display and volunteer EEG recording during stimulus presentation.

### 3.3. Feature Extraction: Training Methodology

The stepwise linear discriminant analysis (SWLDA) as described by Speier et al.[10] and originally used in [3] was used as a classifier. SWLDA is a stepwise method that sequentially adds or removes features based on their statistical significance, optimizing the model for better classification. It was chosen for its efficiency in handling high-dimensional data and its proven effectiveness in BCI applications [39].

SWLDA utilizes a discriminant function, which is determined in the training step [40]. The SWLDA classifier was trained using one of two methods: Across-subject cross-validation (ASCV) (also known as leave-one-subject-out cross-validation) and within-subject cross-validation (WSCV). Although both use SWLDA, note that these two techniques fundamentally differ in how entire subject responses are split between what was used for training and what was used for testing.

a. **ASCV:** Of the *N* subjects, one subject was chosen as the test subject, while the data from the other *N −1* subjects were used to train the SWLDA classifier. The SWLDA classifier then attempted to take the data of the test subject and spell out the target phrase. Note that this technique finally produces one classifier table for all *N* subjects.
b. **WSCV:** In WSCV, a three-fold cross-validation was performed using SWLDA on the EEG data of each individual subject (unlike ASCV), resulting in a unique classifier output for each subject. More specifically, the output phrase contained at least 20 characters and had 120 flashes for each character which was split into three sets. During the first test iteration, the first set was chosen as the test phrase, while the other two sets were used to train the SWLDA classifier. On completion, the same procedure would repeat, but with the second and third sets being the test phrases during the second and third trial, respectively. This method would then be replicated on all subjects.

During training, class labels were predicted using ordinary least squares. Next, the forward step-wise analysis adds the most important features, and the backward analysis step determines and removes the least important features. This process repeats until the number of features remains constant after a set number of iterations or until the number of required features is met. The final selected features are then added to the discriminant function. The flash score *i* for the character *t*, 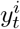can then be calculated as the dot product of the feature weight vector with the features of the signal from that trial.

### 3.4. Virtual flashboard design using a language model

The virtual flashboard schemes used in this study follow [13]. In brief, the highlighted set of characters is modified without changing the underlying flashboard structure, allowing the approach to overlay onto any existing flashboard in current use. Two schemes are evaluated:

a. **Sequential flashboard:** Characters are highlighted in frequency-weighted order rather than alphabetically, so that higher-frequency characters are flashed earlier. See Figure 4 and reference [13] for details.
b. **Diagonal flashboard:** The most likely characters are arranged along diagonals (Figure 5), forcing contextually similar characters to flash separately. The most likely letters *e, t, a, i, n, o* are placed on the main diagonal, with less likely letters on adjacent diagonals.

**Figure 4:**
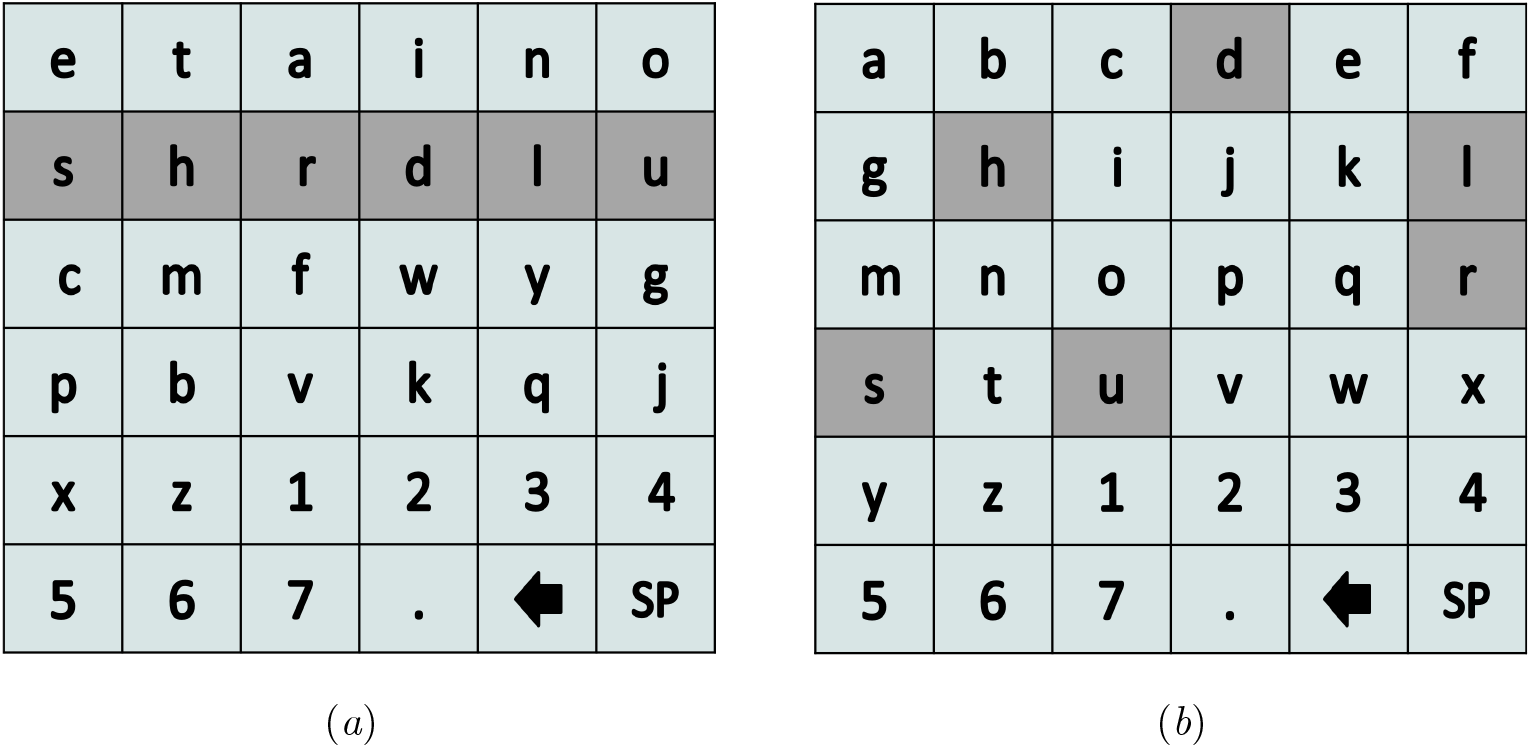
a) Probabilistic flashboard highlighting order (Prob(*Next character/Earlier spelled characters*) in a sequential frequency-sorted flashboard. *SP* in the flashboard denotes space between words b) Sequential flashboard characters shown in Figure 4a are *virtually* mapped onto a conventional alphabetical flashboard

**Figure 5:**
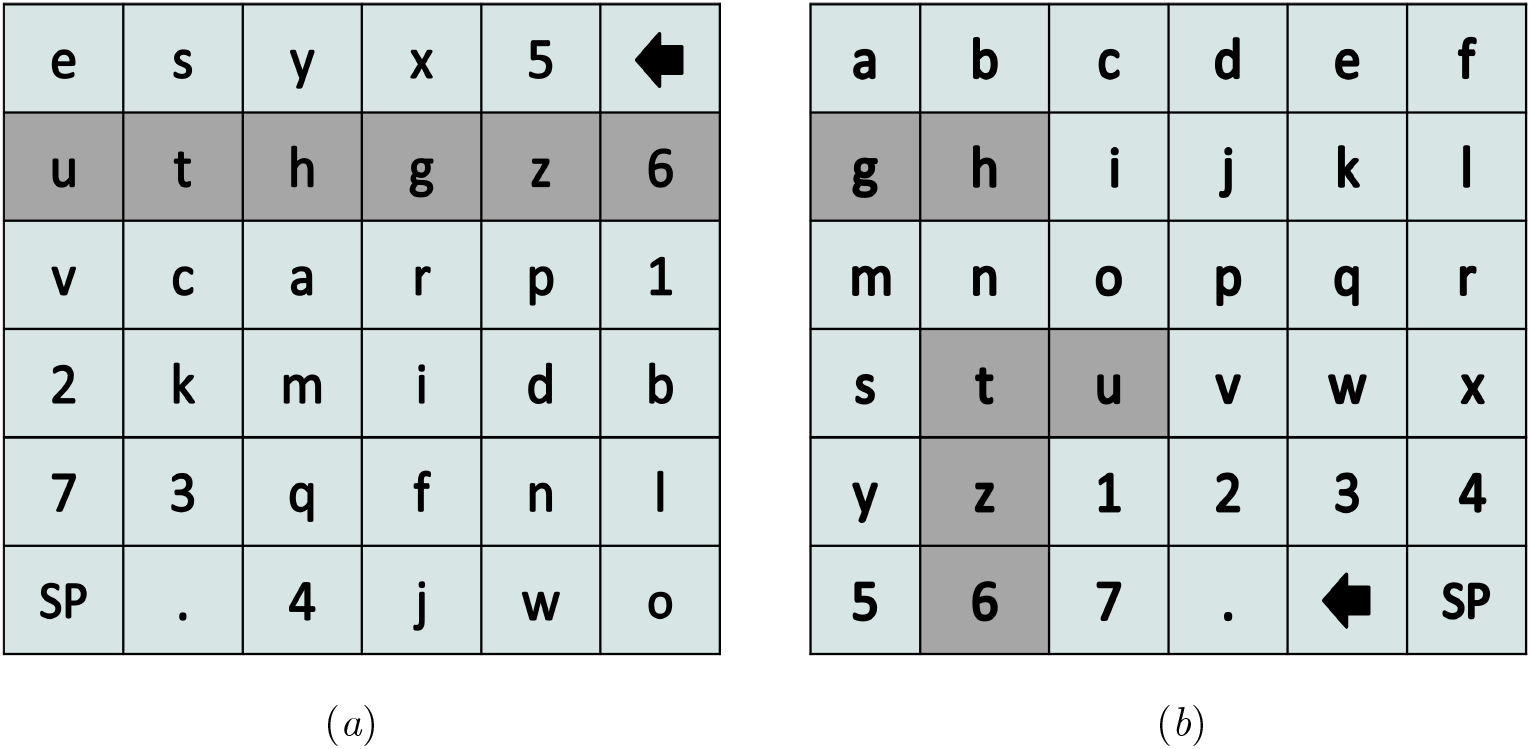
(a) Highlighting in a diagonal flashboard design where the probabilities represent the Prob(*Next character/Earlier spelled characters*). Most likely characters *e, t, a, i* … are organized along diagonals. Note that *SP* in the flashboard denotes space between words b) Diagonal highlighted flashboard characters in Figure 5a are *virtually* mapped onto a conventional alphabetical flashboard

### 3.5. Further optimization of the scanning order

Weighted scanning order and dynamic stopping follow [13]. In brief, weighted scanning highlights character sets in order of probability rather than round-robin. Dynamic stopping terminates the current character flash sequence as soon as a decision threshold is reached, immediately beginning the next character. Dynamic stopping was applied in all simulation conditions reported in Table 1 as well as in the physiological ceiling computation in Section 3.11.

**Table 1:** Information transfer rate (mean ± standard deviation format) in bits/min of various schemes for WSCV and ASCV subject training. The scheme with the highest mean ITR is shown in bold.

| Scheme | WSCV ITR<br>(bits/min) | ASCV ITR<br>(bits/min) |
| --- | --- | --- |
| Random | 51.93 ± 11.2 | 33.95 ± 14.4 |
| Deterministic | 53.42 ± 11.4 | 34.49 ± 14.9 |
| FreqSorted | 53.44 ± 10.8 | 35.44 ± 15.0 |
| Diagonal | 54.43 ± 11.0 | 36.51 ± 15.4 |
| Char Bound | 55.86 ± 10.5 | 37.97 ± 14.6 |
| Dijkstra | 63.59 ± 10.1 | 46.05 ± 17.8 |
| Roberta | 59.01 ± 10.6 | 41.19 ± 16.6 |
| Bert | 62.81 ± 10.1 | 45.25 ± 17.6 |
| XLnet | 65.51 ± 9.8 | 48.18 ± 18.4 |
| Bart | 71.64 ± 8.6 | 55.23 ± 20.1 |
| Bloom | 72.49 ± 9.3 | 56.69 ± 19.7 |
| Opt | 75.04 ± 7.4 | 59.97 ± 20.4 |
| Falcon7 | 74.91 ± 7.3 | 59.89 ± 20.3 |
| MPT_30b | 74.42 ± 7.4 | 59.69 ± 20.3 |
| GPT-NeoX_20b | 74.72 ± 7.4 | 59.7 ± 20.3 |
| GPT-Neo | 74.92 ± 7.4 | **60.22 ± 20.4** |
| GPT-J | **75.36 ± 7.2** | 59.90 ± 20.3 |
| GPT-2 | 75.31 ± 7.5 | 59.75 ± 21.0 |
| LLaMA 3.1 | 75.09 ± 7.2 | 60.18 ± 20.4 |
| LLaMA 3.3 | 75.10 ± 7.2 | 60.17 ± 20.4 |
| Word Bound | 78.21 ± 6.4 | 62.84 ± 21.8 |

### 3.6. Word prediction algorithms

Similarly to word completion suggestions that are available when messaging using smartphones, flashboard efficiency can be likewise increased by replacing unlikely characters with word choices. This section describes algorithms to compute these word choices.

a. **Dijkstra’s Algorithm:** The classic technique for traversing a tree or a graph is either a depth-first search or a breadth-first search, which are well known algorithms [19]. Although computationally efficient, a key limitation is that it does not provide a technique for evaluating OOV words, which occur when the partially spelled word is absent from the language model. In contrast, our approach is to use Dijkstra’s algorithm [18] for word prediction. This is implemented in the form of a trellis similar to dynamic programming techniques as shown in Figure 6. As shown in the figure, every node in a trellis stage has *M* incoming possibilities (where *M* is the number of unique characters on the flashboard) out of which only the one with the highest probability is retained. In doing so, the complexity is reduced, as every possibility does not have to be exhaustively evaluated. Further, as more stages are incorporated into the trellis, longer length word choices are obtained, resulting in higher evaluation time and complexity. For each branch at a given stage, probabilities are determined by using the frequency count of branch characters given prior characters using the language model along with smoothing. At every stage, the highest probability completions then provide the most likely word completions.
b. **Large Language Models:** Numerous early open-source language models were used in this evaluation, such as Roberta [21], BERT [22], XLnet [24], BART [23], OPT [27], GPT-J [28], GPT-Neo [25] and GPT-2 [34],[35]. This was complemented by more recent open-source models such as GPT-NeoX [29], Bloom [26], Falcon [30], MPT [31], LLaMA 3.1 [32] and LLaMA 3.3 [33]. The description provided here will be brief, as the literature on them is easily available in addition to being widely available open-source implementations. Although GPT-4 and GPT-5 have since been released, they are not open-source and do not readily expose word prediction probabilities, precluding their use here. Although modern LLMs are powerful, they cannot reliably predict arbitrary OOV completions without fallback mechanisms. However, they can be combined with Dijkstra’s algorithm (as shown in Figure 7) to account for OOV words, thus creating a layered approach similar in spirit to character-based smoothing algorithms. More specifically, as shown in the figure, in case the GPT-2 word choices result in an empty or partially complete set of choices, one can revert to Dijkstra’s to predict OOV words.

**Figure 6:**
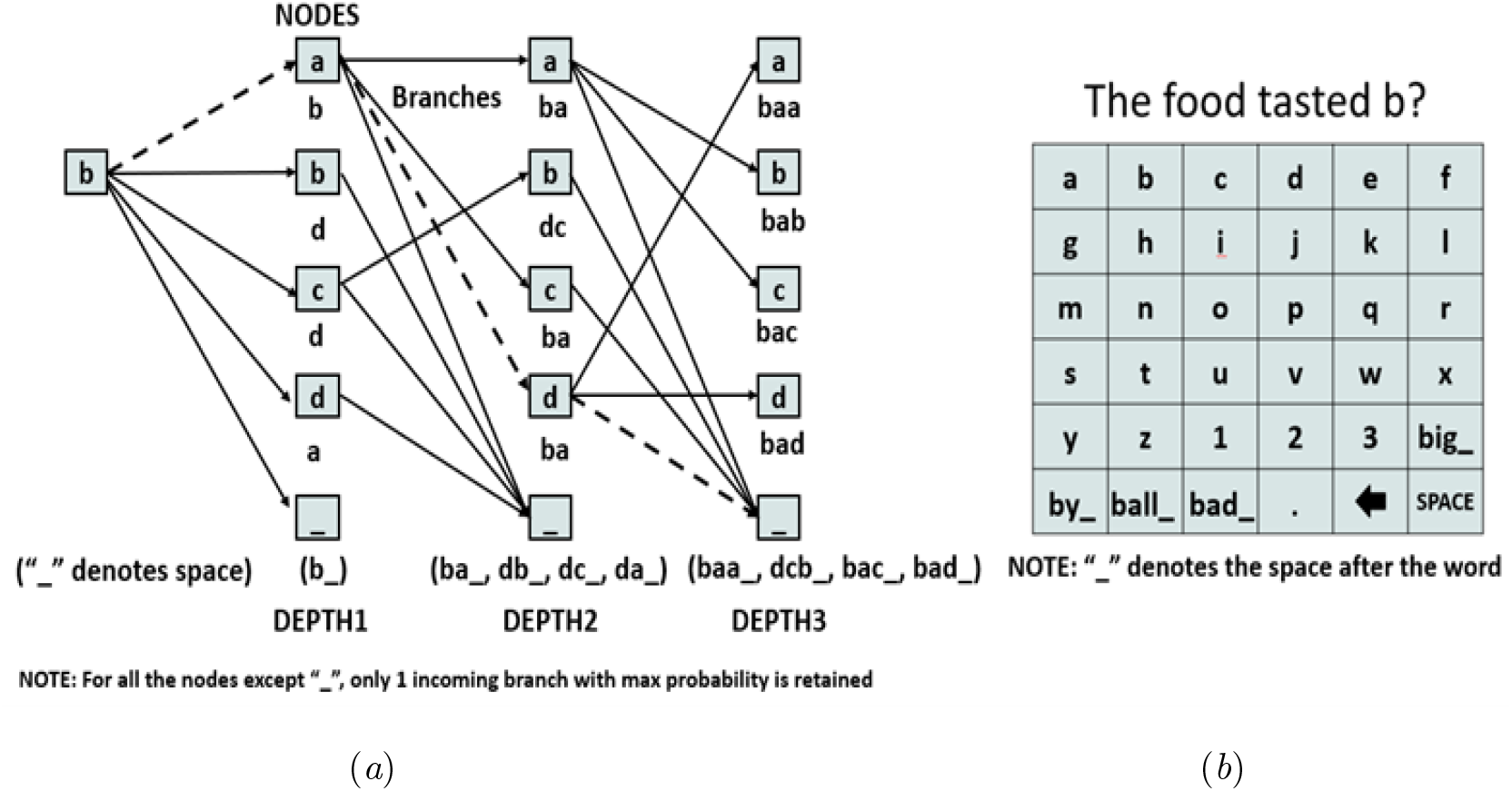
(a) Implementation of Dijkstra’s algorithm to find most likely word completions. (b) Corresponding flashboard showing final most likely word completions with 4 characters have been replaced by the word choices. Note: “ “ represents space between words.

**Figure 7:**
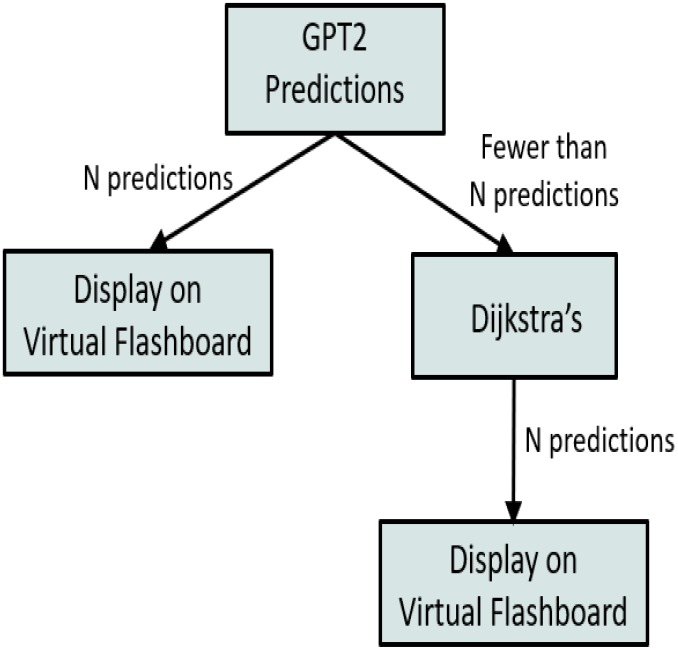
Layered GPT2 word choice selection to obtain most likely *N* word completions with fallback to Dijkstra’s algorithm

### 3.7. Performance Bounds

To quantify how closely practical systems approach optimal performance, we define two theoretical performance bounds: a character bound and a word bound. Each bound is constructed by replacing the language-model prior with an oracle prior that assigns maximum probability to the correct prediction at every decoding step. These bounds are defined within the same decoding framework described in Section 3.8, so the only difference between a practical system and its corresponding bound is the accuracy of the prior.

#### 3.7.1. Character bound

At each decoding step, the oracle assigns a prior probability of 1 to the correct next character and 0 to all other characters. Formally, let *x*^*\**^ denote the correct next character. The oracle prior is then

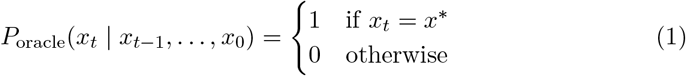

This prior is substituted directly into the posterior in Equations 4–5 in place of the language model prior. Because the prior concentrates all probability mass on the correct character, the decoder requires fewer flash sequences to accumulate sufficient evidence to cross the decision threshold *P*_Thresh_, thus minimizing the selection time. The resulting ITR constitutes an upper bound on what any character-level language model can achieve within this decoding framework, regardless of model architecture or complexity.

#### 3.7.2. Word bound

The word bound is defined analogously at the word level. At each word boundary, the oracle assigns a prior probability of 1 to the correct next complete word among the *K≤* 6 word suggestions displayed on the flashboard, and zero probability to all other word candidates:

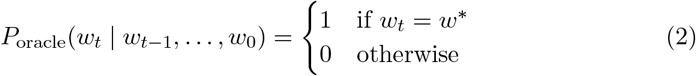

When the correct word appears among the *K* suggestions, selection requires only a single confirmation flash sequence, yielding the minimum achievable selection time for word-level decoding. When the correct word is absent from the suggestion set—as occurs with out-of-vocabulary (OOV) words—the system falls back to character-level decoding using the character oracle prior of Equation (1), ensuring the bound remains well-defined across all tokens in the test corpus.

#### 3.7.3. Validity as upper bounds

These bounds are valid upper bounds for the following reason: the oracle prior eliminates all uncertainty attributable to the language model while leaving the neural decoding process, governed by the EEG signal quality and the SWLDA classifier, unchanged. Any practical language model can at best approximate this oracle, so no real system operating within this framework can exceed the corresponding bound. Crucially, the bounds are subject-specific: because ITR depends on individual P300 signal strength, captured in the attended and non-attended distributions 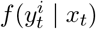 of Equation (3), the bound for each subject reflects that subject’s neural signal fidelity. The bound values reported in Table 1 represent averages across all 78 subjects, consistent with how practical scheme ITRs are reported throughout. It is important to note that these bounds are defined within the constraints of the decoding framework used in this study and do not represent information-theoretic capacity limits of the P300 communication channel. Rather, they quantify the maximum achievable performance under ideal language-model priors while preserving the same neural decoding assumptions. As such, they should be interpreted as practical upper bounds for systems operating within this architecture.

#### 3.7.4. What the bounds capture and do not capture

The character and word bounds isolate the contribution of the language model to overall system performance. The gap between a practical scheme and its corresponding bound reflects residual language modeling error: that is, cases in which the model fails to assign sufficiently high probability to the correct prediction. The gap between the word bound and the physiological ceiling derived in Section 3.11 reflects limitations in neural signal decoding that no language model improvement can address. Together, these two bounds partition the total system performance gap into a language modeling component and a neural decoding component, providing a principled basis for identifying where future research effort is most likely to yield meaningful returns.

### 3.8. Decoder

In this paper, a simple decoding scheme based on thresholding is used to deduce the character or word. First, the score for each flash is computed by 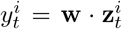 where the weight vector of the feature is determined by extracting the feature as described in the previous subsection. With the assumption that the distributions were Gaussian [10], the *‘attended’* and *‘non-attended’* signals (note that this terminology refers to highlighted and non-highlighted characters on the flashboard) were found and given by the authors.

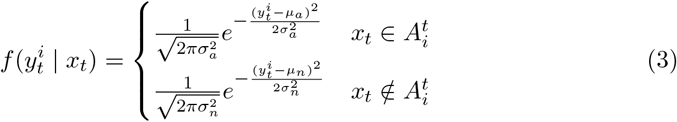

where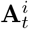 is the set of characters illuminated for the *i*^*th*^ flash for the character *t* in the sequence. *µ*_*a*_, 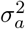, *µ*_*n*_, 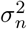are the means and variances of the distributions for the attended and non-attended flashes, respectively.

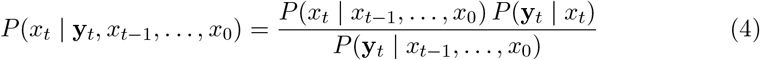

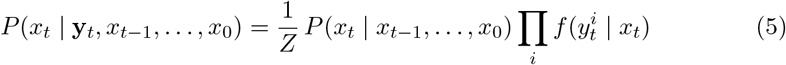

where *P* (*x*_*t*_ *x*_*t−*1_, …, *x*_0_) is the prior probability of a character given the history,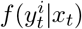 are the probability density functions from equation 3, and *Z* is a normalizing constant. The language model is used to derive the prior as per

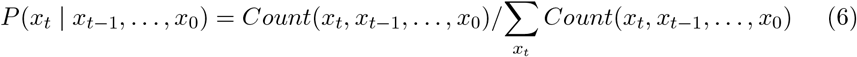

where *Count*(*x*_*t*_, *x*_*t−*1_, …, *x*_0_) is the number of occurrences of the string *x*_*t*_*x*_*t−*1_ … *x*_0_ in the corpus. The threshold probability, *P*_*Thresh*_, is then set to determine when a decision should be made. In all our simulations, *P*_Thresh_ was set to 0.95, consistent with the value used in prior work [12], [13]. The program flashes characters until 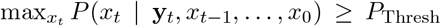or the number of flash sets reaches a pre-determined maximum value. The classifier then selects the character that satisfies arg 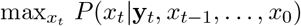.

The subject can correct the typing errors by invoking the backspace provided in the flashboard. As a result, all typing errors are corrected, resulting in zero residual error in the final output text after correction. Alternatively, one can have an auto-correct wherein typed words are checked for presence in a dictionary and erred words are replaced by closest most likely words. This technique is faster, but some mistyped words are likely to either be mis-corrected or left uncorrected, which would result in a non-zero final error rate. Although a vast array of more powerful decoding schemes can be used that involve hidden Markov models, machine learning, and particle filters [10], note that these enhancements only complement the techniques introduced in this paper and therefore do not diminish the results.

### 3.9. Simulation overview

Across all subjects in the dataset, the average waveforms found for the attended and non-attended stimulus responses are shown in Figure 8. Using the EEG signal for the online sessions, the stimulus responses were grouped according to whether they corresponded to the character the subject was currently trying to spell. For each subject, the average response for attended and non-attended stimuli was produced. Then a global average was produced by averaging these signals across subjects. For this discussion, we focus on four channels that we previously determined to be the most influential in our P300 analyses [38]. In each of these channels, a defined negative peak was observed around 200 ms after attended stimuli, followed by a positive peak around 300 ms. The peaks in the Cz channel were slightly later than the peaks in the parieto-occipital channels. In each of these channels, the difference between the attended and non-attended signals was significant in each of these peaks. In addition, there were smaller peaks in both attended and non-attended signals with an interval of 125 ms, which reflects the stimulus onset asynchrony. These peaks likely reflect attended stimuli before or after the current stimulus due to the overlapping response intervals. Statistical variation from such small data sets would limit the confidence in the simulation results. As a result, a large and meaningful data set is used as simulation data. The text of *the Declaration of Independence* (DoI) is used as simulation data [41]. The full length of this document is 7,892 characters after removing all punctuation and special characters. The DoI was chosen as the simulation dataset as it is concise, well-known, and contains words of varying length, including a number of out-of-vocabulary (OOV) words that stress the simulation. Furthermore, to keep the simulation accurate, numerous subject responses are used to provide a wide range of brain activity.

**Figure 8:**
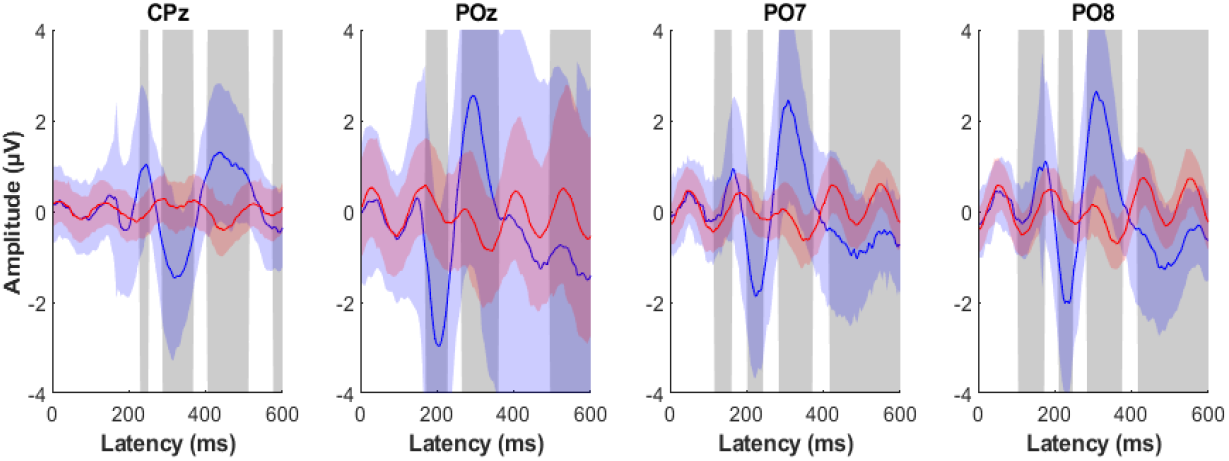
Waveform analysis across all subjects in the dataset, subjects for four channels. These are the average ERPs for target (blue) and non-target (red) stimuli, which are averaged across all dataset subjects. Shaded regions represent the standard error of the mean, and vertical gray regions represent significant differences after using false discovery rate to account for multiple comparisons.

During simulations with the DoI dataset, the attended and non-attended (once again, this terminology refers to whether the desired character is present or absent in the highlighted flashboard character) scores for a given subject were chosen from the laboratory results for that particular subject. The laboratory results are sampled and subsequently modeled using a two-state Markov process representing attended and non-attended responses. This formulation captures the temporal dependence between successive stimulus responses, where the probability of a given state depends on the immediately preceding state, while preserving the binary structure inherent in P300 detection. This approximation provides a tractable way to model correlations across sequential stimuli without introducing significant additional complexity. Higher-order models were not considered to avoid over-parameterization given the temporal resolution of the EEG signals.

For the two-state Markov model, there are four possible sampling combinations depending on whether the current and previous state was attended or non-attended. When a character or word is decoded, it is compared to the intended character or word. In the event of an error, a *Backspace* is initiated as the subsequent character. Following this, the deleted character/word is flashed again. The net effect of this is to *undo* the errors and as errors are corrected during decoding, the final output text contains no residual errors. However, note that this could possibly require multiple attempts and a long follow-up time. The follow-up time could be significantly restricted in addition to tolerating residual error, and this evaluation could be a potential future study.

### 3.10. Performance evaluation metrics

Information Transfer Rate (ITR) is a crucial metric in the evaluation of Brain-Computer Interfaces (BCIs), quantifying the speed and precision of communication [4]. The metric is attractive for several reasons: It is derived from the principles of information theory, it combines competing statistics of speed and accuracy, and it reduces performance to an information transfer problem that can be compared between applications [42]. Following McFarland et al. [4], it is calculated as:

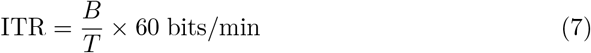

where *B* is the information per selection in bits, defined as:

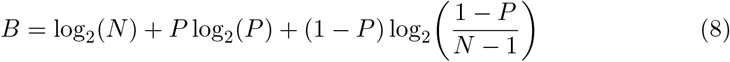

where *N* is the number of characters in the speller (grid size), *P* is the probability of a correct selection, and *T* is the average time per selection, including all pauses and correction overhead. At perfect accuracy (*P* = 1), *B* reduces to log_2_(*N*).

It is possible for the system to generate errors faster than the user can correct them. However, in our analysis, in almost all cases, we see high enough accuracies that users can make all corrections as they happen, resulting in a perfect output. This represents an upper-bound evaluation scenario. In practical deployments, systems may allow a small residual error rate in exchange for improved communication speed, resulting in a different operating point on the speed-accuracy trade-off. To allow for finite duration simulation of a particular character or word, when the simulation requires more than 75 flashboard scans, it is abandoned, and an ITR of 0 (100% error rate) is assigned for that particular character or word and the simulation then moves on to the next character or word.

Publications in this area [46],[47] have used sensitivity and specificity as performance metrics. However, metrics such as sensitivity and specificity measure the frequency of correct letters in a subject’s output. In the current study, all errors are manually corrected, so no errors remain in the output text. Therefore, metrics such as sensitivity and specificity are 100% for all subjects by necessity.

Some errors (e.g. obvious typos) do not affect the readability of text and could therefore be allowed to remain in the final output without affecting the meaning. Allowing these errors to remain could then potentially result in a more efficient system because it would not waste time forcing the user to make these corrections. Language models could also be used to automatically correct some of these errors [10],[11] if they are not manually corrected.

Both the Shapiro–Wilk and Kolmogorov–Smirnov tests [44] reject normality (*p <* 10^*−*3^). Consequently, nonparametric methods are used for hypothesis testing. The Kruskal–Wallis test [44] indicates a significant difference across all schemes (*p <* 10^*−*6^), rejecting the null hypothesis that the ITR and error rates originate from the same distribution. Pairwise Wilcoxon signed-rank tests further demonstrate that the diagonal scheme (*p <* 0.001) and GPT-2 (*p <* 0.001) each significantly outperform the Random baseline. Convergence among LLM architectures is therefore assessed descriptively using Table 1.

### 3.11. Theoretical Performance Bound and Shannon Limit

To understand the significance of this convergence, it is instructive to place the word performance bound within a broader information-theoretic context. This allows the remaining performance gap to be attributed to specific system components and provides a principled basis for identifying where future improvements are most likely to be productive.

#### 3.11.1. Information Content per Selection

The general ITR formula defined in Equations (7)–(8) reduces, at perfect accuracy (*P* = 1), to:

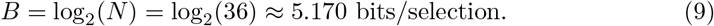

This quantity is fixed by the grid size and is largely independent of the language model or decoding strategy used.

#### 3.11.2. Physiological Minimum Selection Time

The minimum selection time is constrained by P300 response physiology. With an SOA of 125 ms and 12 flash sets per sequence, the selection time assuming a single flash sequence with dynamic stopping is:

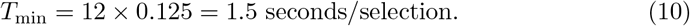

This represents an optimistic lower bound on selection time, assuming reliable P300 detection from a single flash sequence. In practice, multiple sequences are typically required for reliable detection. A more physiologically conservative estimate, assuming a minimum of two flash sequences, yields:

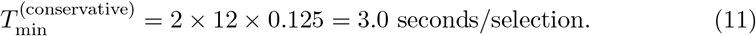

#### 3.11.3. Maximum Theoretical ITR

Substituting into the ITR formula gives:

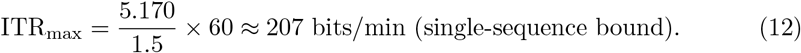

The single-sequence estimate of 207 bits/min represents an idealized upper bound under single-sequence detection assumptions, as reliable P300 detection from a single flash sequence is not typically achievable in practice. Under the conservative two-sequence assumption:

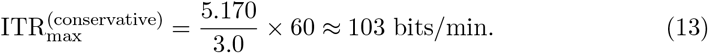

The achievable physiological ceiling therefore lies in the range of approximately 103–207 bits/min.

### Ethics statement

This study was conducted according to the principles embodied in the Declaration of Helsinki and according to local statutory requirements. The study was approved by the UCLA Institutional Review Board (IRB #11-002062; initial approval 21 June 2011, continuously active). All participants provided written informed consent prior to participation. Subject recruitment was carried out between 21 June 2011 and 1 January 2021.

## 4. Results

This section details the performance evaluation of various schemes. Figure 9 shows the performance of different schemes for the 78 subjects in a *violin* plot along with an embedded traditional box and whiskers plot. The violin plot depicts the density of the data in the form of a histogram, while the embedded box and whiskers plot provides relevant statistics. In this case, the classifier was individually trained on each subject (*WSCV* training of subsection 3.3). A maximum of six word suggestions were provided, with numbers in the flashboard replaced by word suggestions. Note that the final error rate in the output text is zero in all these cases. To clarify, the ITR was calculated as defined in Section 3.10.

**Figure 9:**
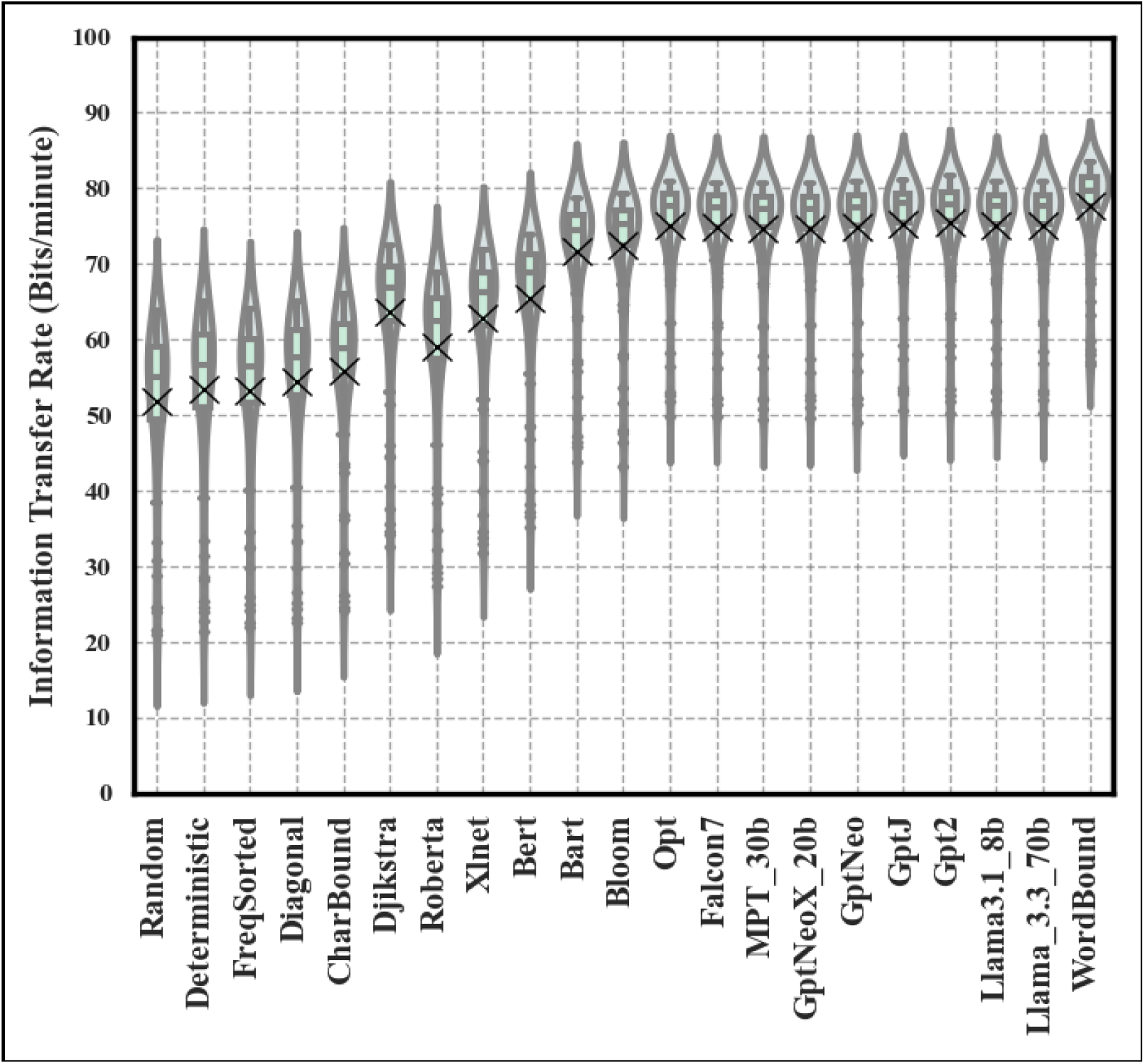
ITR performance comparison of different schemes for the *“Within Subjects”* (WSCV) training **NOTE:** Violin plot containing an embedded box-and-whiskers that shows the minimum, first quartile, median, third quartile, and maximum data. *‘X’* is the mean ITR.

Table 1 shows that GPT-J based word completion achieved the highest average WSCV ITR (75.36 bits/min), with GPT-2, LLaMA, Falcon within 1–2% of it. WSCV schemes consistently produced lower variance than ASCV training. For LLaMA, the WSCV standard deviation was 7.2 bits/min, approximately one third of the ASCV value of 20.4 bits/min. Note that the width of each of the violin plots indicates the ITR standard deviation. In ASCV training, similar performance trends are seen with the GPT-Neo configuration, which provided the highest ITR (60.22 bits/min), as shown in Figure 10 and Table 1. As with WSCV, most models fall within 1–2% of peak ASCV performance. Figure 10 shows the ASCV equivalent of Figure 9.

**Figure 10:**
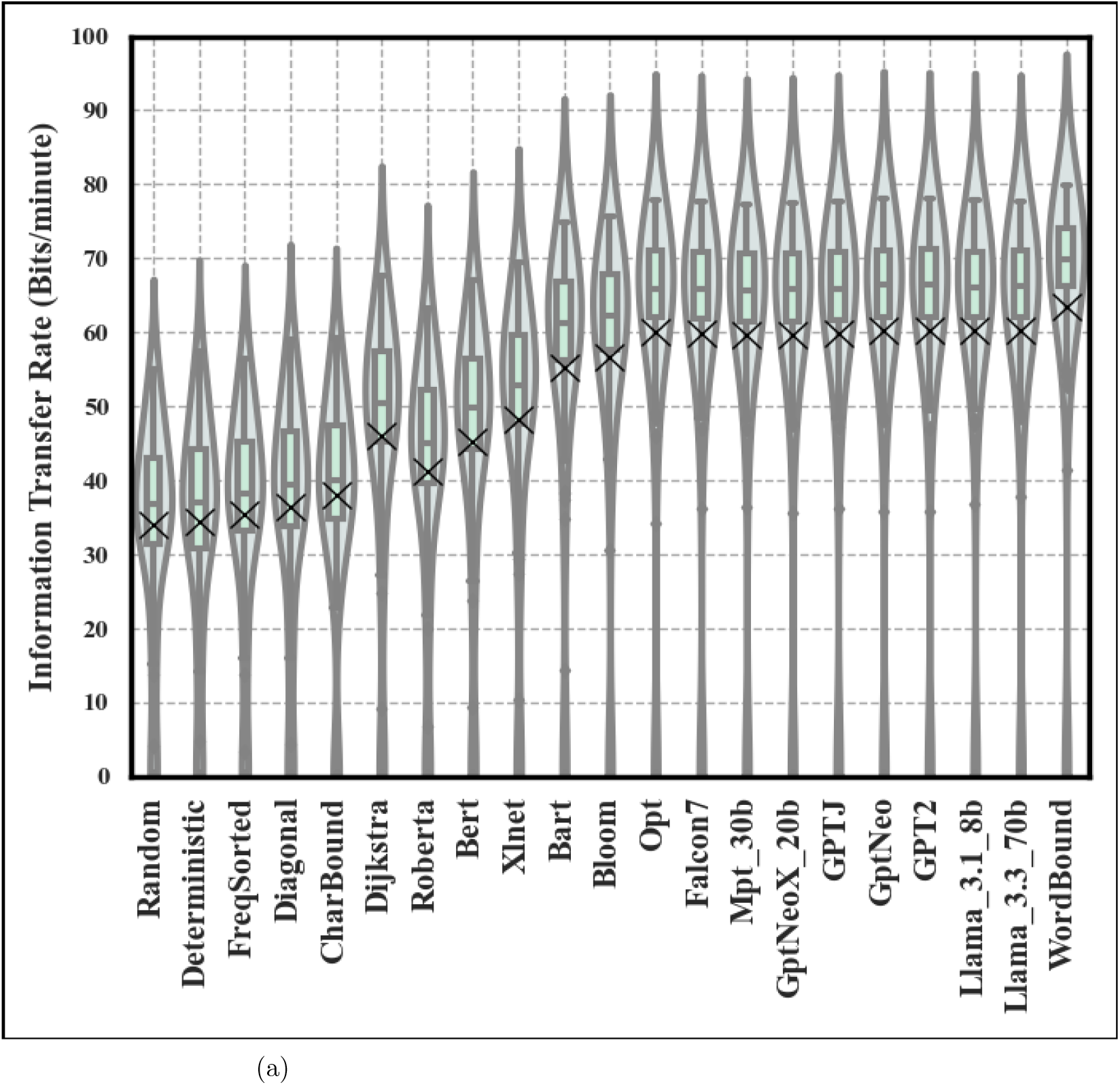
ITR performance comparison of different schemes for the *“Across Subjects”* (ASCV) training

In Figures 9, 10 and Table 1, the character and word performance bounds computed for each subject (as described in Section 3.7) are also shown. These limits are a means of studying the efficacy of each particular scheme compared with the best achievable. The performance of the diagonal scheme is within 2% of the character bound as shown in Table 1 (54.43 vs. 55.86 bits/min for WSCV and 36.51 vs. 37.97 bits/min for ASCV). Following similar trends, regardless of the WSCV/ASCV training methodology, GPT-2/GPT-Neo/LLaMA/MPT/Falcon achieve performance to within 5% of the word bound (for example, Falcon7 is 74.91 vs 78.21 bits/min for WSCV and 59.89 vs 62.84 bits/min for ASCV).

Figure 11 shows the performance of traditional schemes augmented with effective word completion and prediction mechanisms. As shown in Figure 4, by employing the virtual flashboard concept, LLM-based word completion schemes can act as a simple overlay on conventional flashboards while achieving near-optimal performance with respect to the language-model-driven decoding bound.

**Figure 11:**
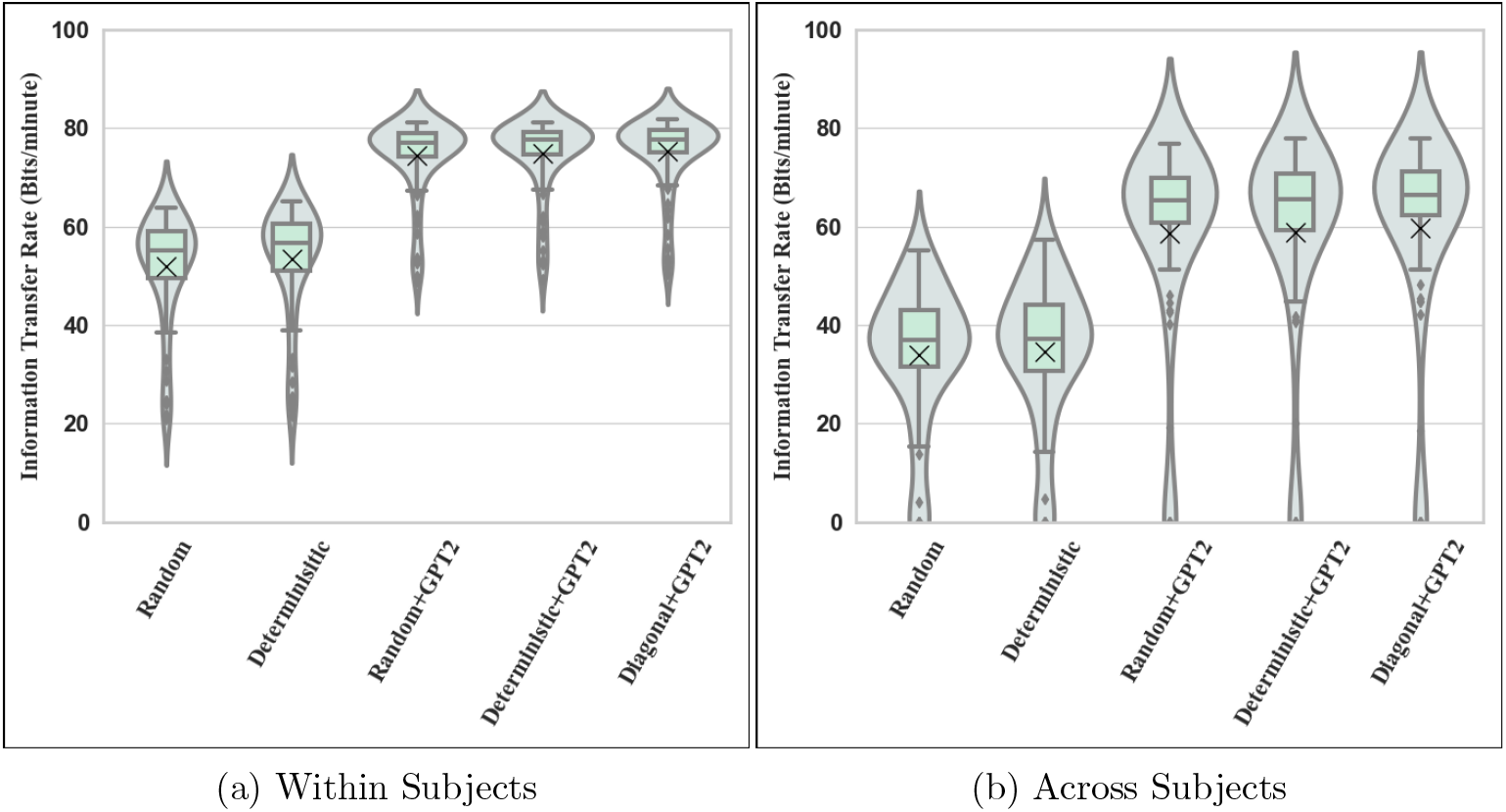
Violin plot of the information transfer rate of conventional flashboards, showing the effect of word completion on performance improvement for *a) “Within Subjects”* (WSCV) and *b) “Across Subjects”* (ASCV). **NOTE:***”Regular”* refers to the traditional random scan with alphabetical flashboard while *“Regular-Wcomp”* refers to the same but with word suggestions included

## 5. Discussion

The central finding of this study is that LLM-assisted P300 spellers have largely exhausted the performance gains available from language modeling alone within the current decoding framework. Across all evaluated architectures spanning early models such as BERT and RoBERTa through recent models including LLaMA and Falcon, the ITR values converge within 2–5% of the theoretical word-prediction bound, indicating that further model scaling is unlikely to yield meaningful additional improvements for this application. These simulations were conducted on a standard HP laptop equipped with an Intel i7 processor and 16 GB of memory, and ITR was calculated as defined in Section 3.10.

In the within-subject (WSCV) analysis, the diagonal scheme achieved a mean ITR of 54.4 bits/min, modestly outperforming the random scheme (51.9 bits/min, *p <* 0.001) across all reported metrics. Importantly, as illustrated by the virtual flashboard technique shown in Figures 4 and 5, the underlying flashboard structure remains unchanged; it is the highlighting of the characters that varies with the particular scheme used. Hence, this is expected to have negligible impact on system complexity and user attention.

In Figures 9 and 10, when the Dijkstra algorithm is used for word completion, the ITR gain increases to 63.6 bits/min, representing an increase of almost 15%. When GPT-2 is used in conjunction with Dijkstra’s algorithm, the ITR further improves to 75.3 bits/min (*p <* 0.001), resulting in a net improvement of about 28%. This value falls within 4% of the word performance bound, indicating limited residual gain from further improvements in the language model. In addition, LLM use also substantially reduced the retry rate. To further increase ITR, one could potentially explore predicting not only the next word, but also performing multi-word predictions. However, regardless of the specific algorithm used, word completion algorithms provide substantial gains over schemes that rely solely on character prediction, consistent with previous work [6]. It is worth noting that by applying smoothing, flashboard character probabilities for 17 biwords in the DoI dataset that were out of vocabulary (OOV) from the biword language model were obtained by transitioning to the word and trigram models.

Across subjects (ASCV), performance (Figure 10) shows a trend similar to WSCV. However, compared to WSCV, more subjects exhibit lower ITRs as shown in Figure 10. The diagonal character flashboard showed a 7.6% improvement over the regular flashboard with random flashing, while Dijkstra’s and GPT-2 word completion provide 35.7% and 76% gain, respectively. Overall, these results demonstrate that the gains generalize across both within-subject and across-subject classifiers. This is particularly important for practical deployment, as across-subject classifiers significantly reduce or eliminate the need for subject-specific calibration.

A key finding is that performance gains are largely independent of the specific architecture of the model. The results presented here indicate that a wide range of models evaluated (including GPT-2, LLaMA, Falcon, and others) achieve nearly identical performance. In both within-subject and across-subject evaluations, the variation between models is typically within 1–2%, suggesting convergence across architectures.

The theoretical ceiling complements the word prediction bound introduced in Section 3.7, which captures the limits of language-model-driven decoding. Together, they allow the total system-level performance gap to be partitioned into a language modeling component and a neural decoding component.

As shown in Table 2, the word performance bound of 78.21 bits/min corresponds to approximately 76% of the conservative theoretical ceiling and roughly 38% of the idealized single-sequence ceiling. The gap between the word bound and the physiological ceiling, approximately 25–129 bits/min, cannot be recovered through improvements in language modeling alone. This remaining gap is primarily governed by the number of flash sequences required for reliable P300 detection, as well as the fidelity of EEG signal acquisition and the efficiency of neural decoding. Improvements in classifier design and signal processing therefore represent the most promising avenues for further performance gains beyond what language modeling can provide.

**Table 2:** Comparison of WSCV ITR values against the theoretical Shannon ceiling derived from McFarland et al. [4]. Percentages are relative to the conservative ceiling (*∼*103 bits/min), assuming a minimum of two flash sequences per selection.

| **Scheme** | **ITR (bits/min)** | **% of Conservative Ceiling** |
| --- | --- | --- |
| Random (baseline) | 51.93 | 50% |
| Diagonal scheme | 54.43 | 53% |
| Best LLM (GPT-J, WSCV) | 75.36 | 73% |
| Word bound (WSCV) | 78.21 | 76% |
| Conservative ceiling | $\sim 103$ | 100% |
| Single-sequence ceiling | $\sim 207$ | — |

These findings represent a shift in understanding compared to prior work [13]. Rather than viewing language models as a continuously improvable component driving system performance, our results indicate that the primary bottleneck may now lie in other aspects of the system, such as neural signal quality, feature extraction, or decoding strategies. In this context, improvements in EEG acquisition, signal processing, and adaptive decoding may provide greater benefits than further scaling of these models.

Retrofitting, which refers to the addition of word prediction to standard speller schemes, enables performance to approach near-optimal levels in both within-subject and across-subject analyses (Figure 11). In particular, these gains are largely independent of the specific model used, further supporting the observation that these models exhibit convergent performance. This result suggests that existing speller systems can be significantly enhanced through the integration of language-model-based decoding without requiring fundamental changes to their structure. More importantly, it reinforces the conclusion that contextual word prediction plays a dominant role in achieving near-optimal performance, whereas further gains from increasingly complex models are likely to be limited.

A primary limitation of this study is the reliance on offline simulation, which assumes that neural signals are not influenced by real-time feedback and user adaptation. In practical BCI use, factors such as fatigue, attention, and feedback-driven behavioral changes can affect performance and may reduce achievable ITR compared to simulation. In addition, dynamic stopping behavior may differ under real-time conditions. While prior work has shown consistency between offline and online trends [10, 11], the results presented here may overestimate absolute real-time performance. Accordingly, these findings should be interpreted as comparative and upper-bound estimates, with validation in online settings remaining an important direction for future work.

Furthermore, the use of simulation enables evaluation over long text passages containing rare and out-of-vocabulary (OOV) words, which are difficult to study in controlled online experiments where subjects typically generate shorter, more common phrases. As a result, the present study focuses on comparative evaluation of language model performance across system configurations, rather than establishing absolute real-time ITR benchmarks.

Another limitation concerns generalizability to the target clinical population. All subjects in this study were healthy adults aged 20–35, whereas individuals with ALS often exhibit reduced P300 amplitudes, increased latency, and greater signal variability due to disease progression and fatigue. As a result, the ITR gains observed here may not directly translate to clinical populations. Evaluating the proposed methods in ALS patients under online conditions remains an important direction for future work.

GPT-2 [34] has well-documented contextual limitations, and language acquisition involves more than word prediction alone. However, advances in GPT-based language models, such as GPT-3 [45] and subsequent models, mitigate these limitations, further enhancing their value. Thus, we do not view this as a fundamental limitation of the results in this paper; rather, it opens avenues for experiments with new models that are more context-sensitive.

Overall, the gap between theoretical and practical P300 speller performance has substantially narrowed, shifting the primary bottleneck from algorithmic design to physiological and hardware constraints.

## 6. Conclusion and future directions

This study investigated the role of large language models (LLMs) in improving the performance of P300 speller brain-computer interfaces, with a particular focus on understanding their fundamental performance limits. By evaluating a wide range of modern LLMs within a unified framework and introducing an idealized model to establish theoretical bounds, we provide a systematic analysis of achievable performance.

Across both training paradigms, the evaluated models substantially improved typing speed, raising ITR from approximately 52 to over 75 bits/min in within-subject settings and from 34 to approximately 60 bits/min in across-subject settings. Importantly, these gains were consistent across multiple model architectures, with most models operating within 2–5% of the theoretical performance limit.

The most consequential finding is not the magnitude of these gains, but their convergence. Despite spanning a wide range of architectures and scales, the evaluated models cluster within 5% of the theoretical word-prediction limit, indicating that further sophistication of the language model is unlikely to produce significant additional benefit for this application. This represents a fundamental shift in where the performance bottleneck lies: the limiting factor in modern P300 spellers is no longer the quality of language modeling, but the fidelity of neural signal acquisition and decoding. Future P300 speller improvements will therefore depend more on advances in neural decoding and signal acquisition than on further language model scaling.

In addition, the use of cross-subject classifiers demonstrates the feasibility of reducing or eliminating subject-specific calibration, which is a key requirement for scalable real-world deployment. The combination of near-optimal language-model-based decoding and calibration-free classification represents a significant step toward practical BCI communication systems.

Future work will focus on validating these findings in online experiments and exploring complementary approaches, such as adaptive thresholding and improved neural decoding techniques, to further enhance system performance.

## Data availability

The code and simulation framework used in this study are available at: https://github.com/nithinparthasarthy/P300Speller.git. The EEG dataset used in this study was collected under IRB #11-002062 approval and is available from the corresponding author upon reasonable request, subject to IRB, institutional and ethical restrictions.

## Appendix A. Kneser–Ney Smoothing Formulation

Note that the convention adopted in the following terminology is that *Model*(‘^*\**^ ‘) denotes the frequency of occurrence of the characters ‘^*\**^ ‘. For example, *biword model(‘the quick’)* is the frequency of the word *‘the quick’* in the biword language model. Also, as an example of the syntax of conditional probability, *p biword*(‘*c*’ |‘*the qui*’) is the probability in the biword model of the character ‘*c*’ given that the partial phrase ‘*the qui*’ has been decoded.

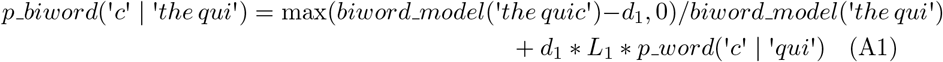

where *L*_1_ is the normalizing constant equal to the number of distinct letters that follow ‘*the qui*’ in the biword model divided by the biword model count of ‘*the qui*’. If *biword model*(‘*the qui*’) = 0, *L*_1_ = 1 to take care of out-of-vocabulary (OOV) conditions.

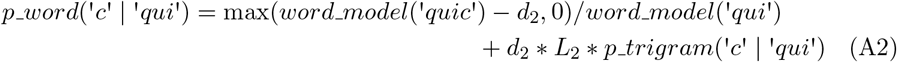

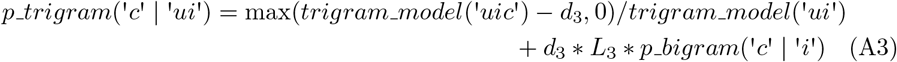

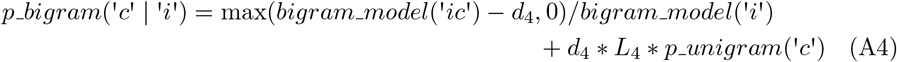

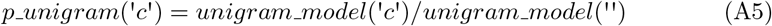

where *unigram model*(‘‘) = *total model entries* and *d*_1_, *d*_2_, *d*_3_, *d*_4_ are tunable parameters with 0 *<*= *d*_1_, *d*_2_, *d*_3_, *d*_4_ *<*= 1. All these *d* values were set to the commonly used midrange default of 0.5. Just as *L*_1_ is a biword model normalizing constant in equation A1 whose derivation was described earlier with an example, *L*_2_, *L*_3_, *L*_4_ are also normalizing constants similarly obtained from the corresponding lower-order language models. More specifically, in the example used, *L*_2_ is the normalizing constant equal to the number of distinct letters that follow ‘*qui*’ in the word model di-vided by the word model count of ‘*qui*’. Furthermore, for satisfying the OOV conditions, if *word model*(‘*qui*’) = 0, *L*_2_ = 1. If *trigram model*(‘*ui*’) = 0, *L*_3_ = 1 and if *bigram model*(‘*i*’) = 0, *L*_4_ = 1 thereby accounting for the values of *L*_1_… *L*_4_ in all OOV cases. In summary, using equations A1-A5, the most complex model in Figure 2 defaults to lower-level models when processing OOV characters.

